# Peptide binding specificity prediction using fine-tuned protein structure prediction networks

**DOI:** 10.1101/2022.07.12.499365

**Authors:** Amir Motmaen, Justas Dauparas, Minkyung Baek, Mohamad H. Abedi, David Baker, Philip Bradley

## Abstract

Peptide binding proteins play key roles in biology, and predicting their binding specificity is a long-standing challenge. While considerable protein structural information is available, the most successful current methods use sequence information alone, in part because it has been a challenge to model the subtle structural changes accompanying sequence substitutions. Protein structure prediction networks such as AlphaFold model sequence-structure relationships very accurately, and we reasoned that if it were possible to specifically train such networks on binding data, more generalizable models could be created. We show that placing a classifier on top of the AlphaFold network and fine-tuning the combined network parameters for both classification and structure prediction accuracy leads to a model with strong generalizable performance on a wide range of Class I and Class II peptide-MHC interactions that approaches the overall performance of the state-of-the-art NetMHCpan sequence-based method. The peptide-MHC optimized model shows excellent performance in distinguishing binding and non-binding peptides to SH3 and PDZ domains. This ability to generalize well beyond the training set far exceeds that of sequence only models, and should be particularly powerful for systems where less experimental data is available.

**Significance statement:** Peptide binding proteins carry out a variety of biological functions in cells and predicting their binding specificity could significantly improve our understanding of molecular pathways. Deep neural networks have achieved high structure prediction accuracy, but are not trained to predict binding specificity. Here we describe an approach to extending such networks to jointly predict protein structure and binding specificity. We incorporate AlphaFold into this approach, and fine-tune its parameters on peptide-MHC Class I and II structural and binding data. The fine-tuned model approaches state-of-the-art classification accuracy on peptide-MHC specificity prediction and generalizes to other peptide-binding systems such as the PDZ and SH3 domains.

## Main text

Sequence-based methods utilize large sets of experimentally validated binding and non-binding peptides to assemble position specific weight matrices or more sophisticated neural networks with several layers to discriminate binder from non-binder peptides (1–7). Methods such as NetMHCpan are the current state-of-the-art to address key biological challenges like MHC peptide binding specificity which is central to the adaptive immune system (T-cell surveillance, differentiation, etc), since they can readily optimize parameters over large sets of binding and non-binding peptides. However, sequence-based methods are limited by their inability to incorporate detailed structural information, and as a result, they have reduced generalizability, particularly in cases where there is less training data. While structure based methods have shown promise to fill this gap, they have been limited by their inability to accurately predict protein and peptide backbone changes which can affect both affinity and specificity, and more importantly, they lack a way to optimize many model parameters on the large amounts of peptide binding data that are often available (8).

We reasoned that the recent advances in protein structure prediction could help overcome both limitations: that of structure based methods in utilizing large amounts of known binding data, and sequence-based methods, in using structural information. AlphaFold (9) and RoseTTAFold (10) predict highly accurate structures (11) and structure quality confidence metrics that have been used to distinguish pairs of proteins which bind from those that don’t with some success (12, 13). However, while these methods can be readily trained with structural data, in their current form it is not straightforward to train on binding data.

We set out to extend these networks to enable simultaneous training on structure and binding data. Because of the importance of the peptide-MHC interaction to adaptive immunity, and the very large datasets available for training and testing, we began by focusing on this binding interaction. We first explored how to provide available sequence and structure data as inputs to AlphaFold to obtain the most accurate peptide-MHC structure models (see Methods), and obtained best performance using single sequence (rather than MSA information) and peptide-MHC structure templates as inputs with the query peptide positionally aligned to the template peptides. This approach modeled peptide-MHC structures with a median peptide backbone RMSD of 0.8 Å for both Class I (**Fig. S1A**) and Class II (**Fig. S1B**) complexes.

We found previously in studies of designed proteins bound to targets (14) that both the AlphaFold residue-residue accuracy estimate, PAE, and the per-residue accuracy estimate pLDDT, are able to partially discriminate binders from non-binders. We carried out predictions for sets of binding and non-binding peptides from (1, 5, 15), and found that both the PAE, calculated between MHC and peptide, and the per-residue accuracy estimate pLDDT averaged over the peptide residues, provided some discrimination of binders from non-binders (**Fig. S2**). However, the discrimination between binders and non-binders was considerably poorer than NetMHCpan, and AlphaFold tended to dock non-binding peptides in the MHC peptide-binding groove (**Fig. 2A**).

We reasoned that the relatively poor discrimination compared to NetMHCpan reflected the lack of training of AlphaFold on negative binding examples generally, and on peptide-MHC binding data in particular. To overcome this limitation, we set out to implement a hybrid structurally-informed classification approach for fine-tuning AlphaFold that incorporates both binder and non-binder peptide-MHC pairs. To enable direct training on binding data, we added a simple logistic regression layer on top of AlphaFold which converts the peptide-MHC PAE values into binding/non-binding predictions. We experimented with different training regimens for optimizing the combined model parameters, and obtained the best results by first fitting the logistic regression parameters, and then subsequently fine-tuning all of the AlphaFold parameters (**Fig. 1**). To maintain structure prediction accuracy, we included peptide-MHC structure prediction examples during training. For cases where the structure was not known, we followed a “distillation” procedure (9) in which AlphaFold models of the complex were used as the ground truth conformation, but with the loss calculated only over the MHC to allow the peptide conformation to adjust during training. Leveraging the wealth of data available on peptide-MHC interactions, we assembled a diverse training set consisting of 10340 peptide-MHC examples, 203 structurally characterized and 5102 modeled peptide-MHC binding examples, and 5035 non-binder examples, distributed across 59 Class I and 58 Class II alleles (see Methods). Training the combined structure prediction-classification model on this set produced consistent drops in the combined loss function (**Fig. S3**), suggesting that the model might be learning to better discriminate binder from non-binder peptides on the basis of structural information. The fine-tuned model developed a tendency to exclude non-binder peptides completely out of the MHC’s peptide-binding groove (**Fig. 2A**).

**Figure 1:**
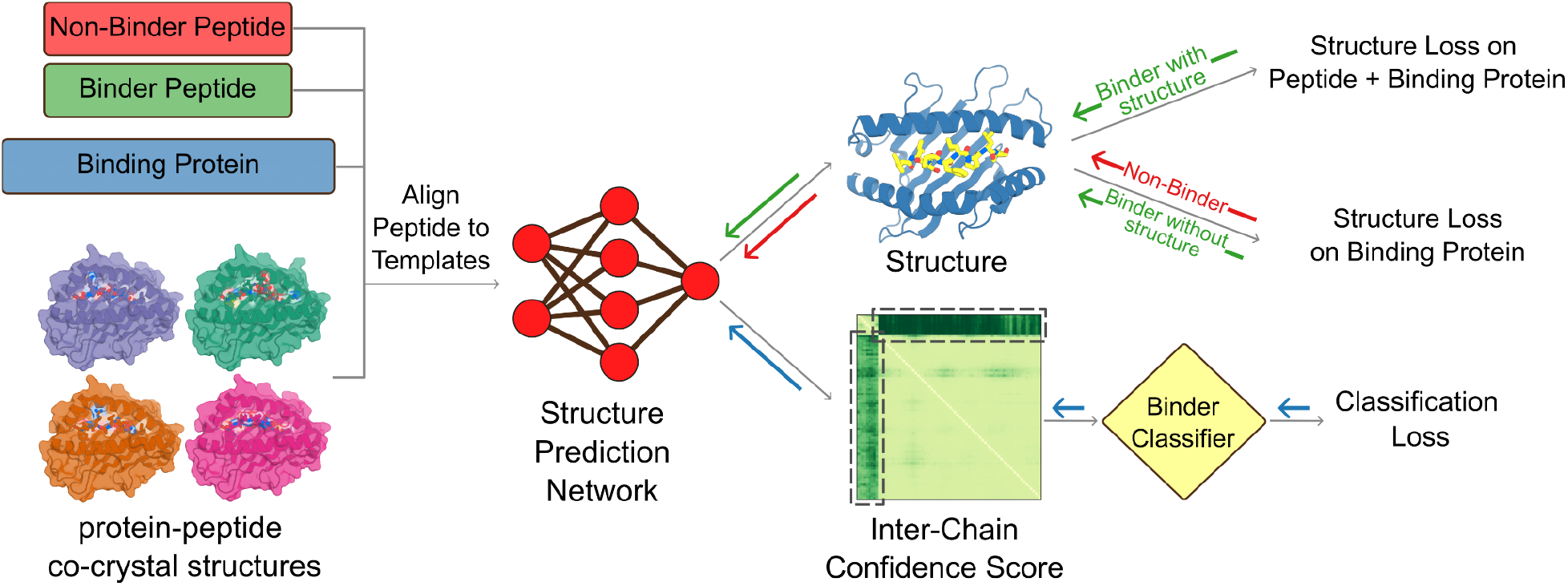
Extending structure prediction networks for binder classification. For fine-tuning the combined model to distinguish binding from non-binding peptides, inputs (left) are sets of known binding (green) and non-binding (red) sequences, the sequence of the target protein(s), and the available co-crystal structures which provide templates for modeling the complex. The peptide sequence is positionally aligned to the peptides in the co-crystal structures, and the structure of the complex is predicted with AlphaFold. A logistic regression classifier converts the output PAE values into a binding/non-binding prediction. During training, the combined model parameters were optimized to minimize the loss terms in the final column, and model weights were updated through gradient backpropagation as indicated by the solid arrows. The classification loss (cross entropy) is computed on all training examples; the structure loss is computed over the entire complex for binding peptides with co-crystal structures, and over the binding protein alone for binding peptides without co-crystal structures and for non-binding peptides.

**Figure 2:**
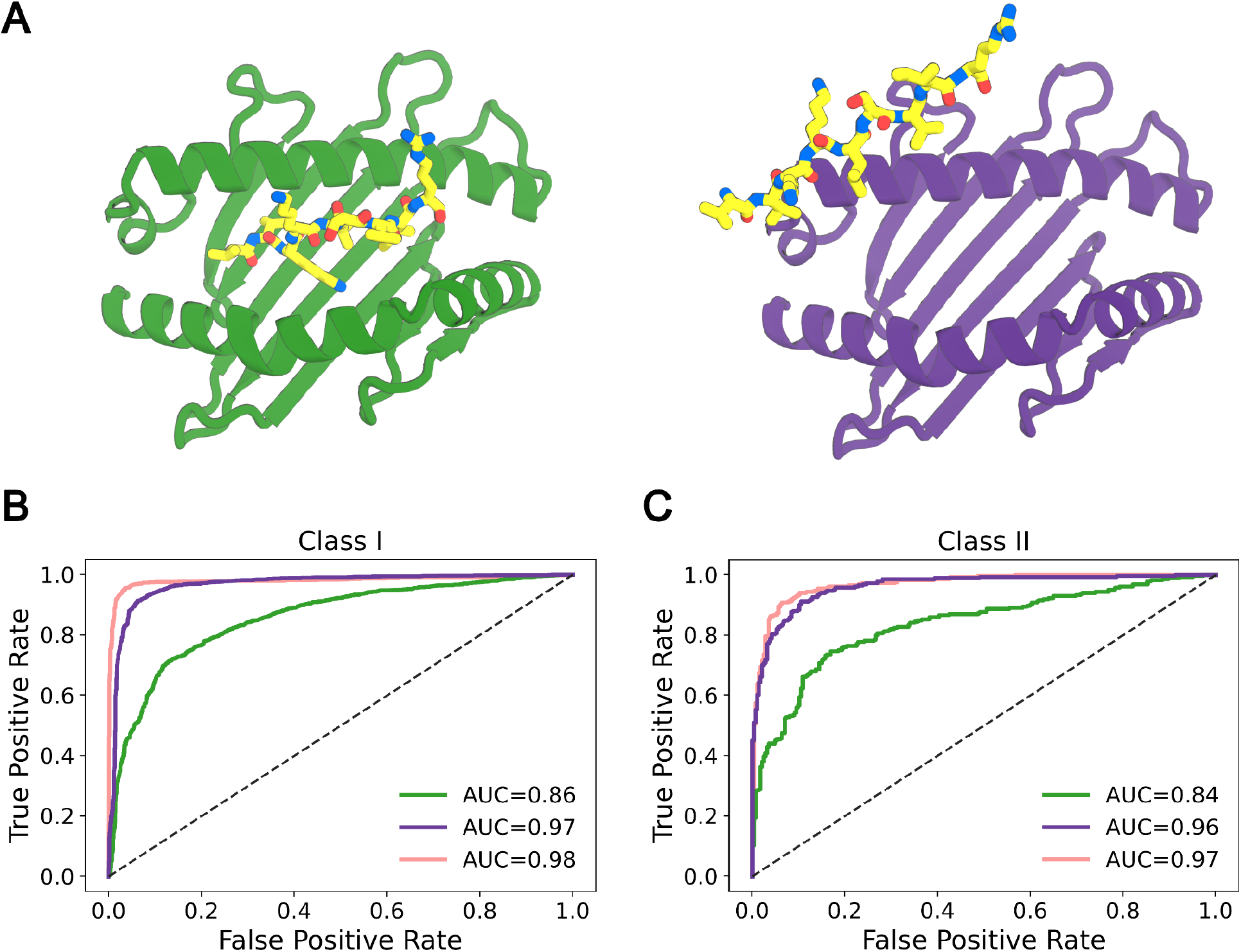
Fine-tuning improves pMHC-peptide binder classification to state-of-the-art levels. **A**. Structure prediction for a non-interacting peptide-MHC pair using AlphaFold before (left) and after (right) fine-tuning on joint structure prediction and binder classification. The peptide is incorrectly predicted to bind the MHC in the first case but not the second. **B-C**. Binder/non-binder peptide classification ROC curves on validation sets (see Methods): (B) MHC Class I; (C), MHC Class II. AlphaFold (green), combined structure prediction-classification model (purple), and NetMHCpan (pink).

We assessed prediction performance on a validation set containing 1641 binder and 1715 non-binder peptide-MHC pairs distributed across 36 Class I and 27 Class II alleles (**Fig. 2**). This set contained no peptide-MHC pairs in common with the training set; results with exclusion of overlapping peptides with the same label (positive or negative) independent of MHC allele are shown in **Figure S4**. During training, the Class I examples were restricted to 9mer peptides, but in the validation set we included Class I peptide-MHC pairs containing peptides of lengths 8, 9, and 10 residues to better assess generalizability of our fine-tuning approach. The combined structure prediction-classification model performs significantly better on this test set than AlphaFold with default parameters (**Fig. 2B, C**). A direct comparison to the state-of-the-art sequence-based predictor NetMHCpan on classical HLA alleles is challenging since the vast majority of available binding data is already contained in its training set. Despite our validation examples being contained within NetMHCpan’s training data, our fine-tuned model performed competitively with NetMHCpan, particularly on Class II, achieving AUROC of 0.97 for Class I (**Fig. 2B**) and 0.96 for Class II (**Fig. 2C**) indicating excellent classification. Our combined prediction-classification approach appears to learn the biochemical space of peptide-MHC binding efficiently rather than relying solely on sequence patterns observed during training.

### Divergence from NetMHCpan on 10mer peptides

Our structure-based approach to peptide-MHC binding prediction is an orthogonal and hence potentially complementary method to purely sequence-based tools such as NetMHCpan. As an illustration, we noticed a striking divergence between structural predictions and NetMHCpan predictions in a scan of the Hepatitis C virus (HCV) genome polyprotein for 10mer peptides predicted to bind to the HLA-A*02:01 allele. Surprisingly, the two top-ranked NetMHCpan peptides (AKLMPQLPGI and ADLMGYIPLV) both had charged amino acids at the first anchor position (peptide position 2, a position which strongly prefers hydrophobic amino acids in known HLA-A*02:01 binders; **Fig. 3B**). By contrast, these two peptides are not predicted to bind to HLA-A*02:01 using our joint structure and binding prediction model (**Fig. 3A**). When predicting affinities for class I peptides of length greater than 9, NetMHCpan aligns target peptides to a 9mer binding model, allowing consecutive positions to “gap out” to match the 9mer model length. For both of these peptides, peptide position 2 was not aligned to the 9mer model, making position 3 the anchor position for the purposes of scoring. This likely explains the highly favorable scores assigned to these peptides, as position 3 is a leucine (a strongly preferred anchor amino acid) in both. To assess more broadly the relationship between the NetMHCpan gap position and the consistency with our predictions, we assigned a rank error equal to the absolute difference in ranks assigned by NetMHCpan and our structural approach to each 10mer peptide window in the target and grouped these rank error values based on the NetMHCpan gap position. As shown in **Figure 3A**, the positions with highest mean rank error are the two canonical HLA-A*02:01 anchor positions (2 and 10, for a 10mer peptide). These results suggest that there may be differences in the reliability of NetMHCpan predictions for peptides of length greater than 9, depending on the location of the gap position(s), and more generally, that our combined structure prediction-classification model may be more robust than purely sequence-based models to the inevitable biological variations in peptide length and other properties.

**Figure 3.**
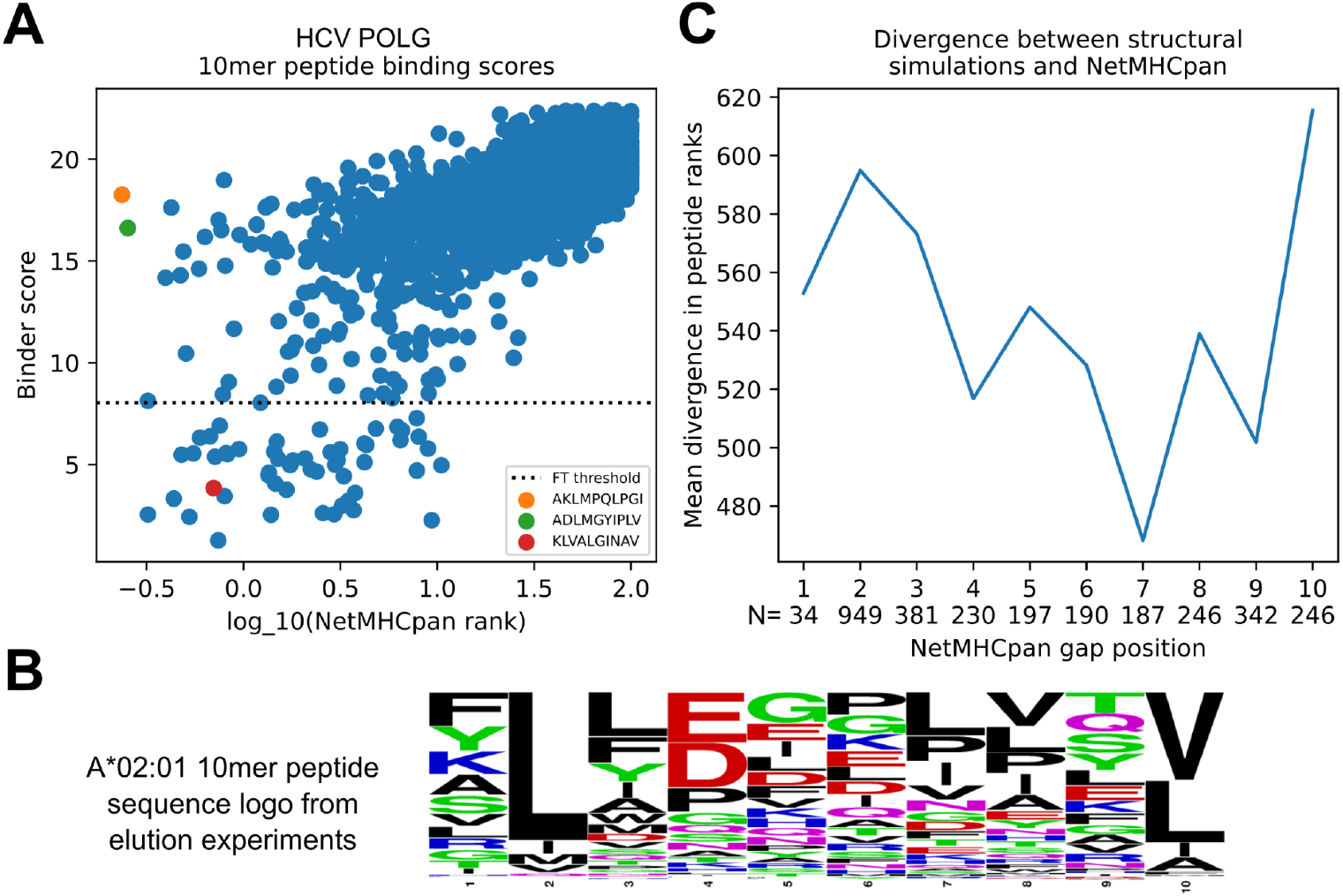
Prediction of 10mer peptide binding highlights differences between sequence- and structure-based approaches. (A) Scatterplot of NetMHCpan rank score (x-axis) versus combined structure prediction-classification model binder score (y-axis) reveals peptides with divergent predictions (lower values correspond to stronger binding). The top two NetMHCpan predicted peptides (orange and green markers) have charged amino acids at the first anchor position (peptide position 2). A known immunodominant T cell epitope (red marker) scores well by both methods. (B) Sequence logo built from 10mer peptides eluted from A*02:01 (5) ^5^ shows strong preference for hydrophobic amino acids at the anchor positions. (C) Average disagreement in peptide rank between NetMHCpan and the structural model (y-axis) is highest when the NetMHCpan gap position is located at one of the two anchor positions (2 and 10).

### Combined structure prediction-classification model extends to other protein-peptide systems

To evaluate the generalizability of our combined structure prediction-classification model beyond the peptide-MHC binding data it was trained on, we assessed performance on two additional systems: peptide recognition by PDZ domains, which bind C-terminal peptides, and peptide recognition by SH3 domains, which recognize proline-rich peptides. The PDZ dataset was taken from Ref. (16) and consists of 17 PDZ domains with an experimentally determined co-complex structure and peptide binders from phage-display selection experiments (17) (**Table S1**). The SH3 dataset consists of 19 SH3 domains with an experimentally determined co-complex structure and peptide binders from phage-display available in the PRM-DB database (http://www.prm-db.org/) (18) (**Table S2**). For comparison with experiments, we performed an “*in silico* selection” experiment in which 20,000 random peptides were modeled and ranked by our structure prediction-classification model (**Fig. 4A**). Position weight matrices (PWMs) were constructed from the top-ranked binding sequences, and these PWMs were compared with PWMs constructed from the experimentally selected binders using two PWM divergence measures proposed by Smith and Kortemme (16). The fraction of top-ranked sequences used to build the PWM (*f*_*PWM*_) is a free parameter in this approach; we varied this parameter from 10^−4^ (i.e, only 2 sequences used to build the PWM) to 1 (all sequences used) and evaluated the accuracy of the resulting PWMs. The prediction errors, plotted as a function of *f*_*PWM*_, are shown in **Figure 4B** for our fine-tuned parameters (purple lines) and for AlphaFold with default parameters (green lines). From these error curves we can see that there appears to be an optimal value for *f*_*PWM*_ somewhere around 0.01: using too few top-ranked sequences likely introduces noise into the PWMs, while using too many of the randomly sampled sequences reduces the degree of enrichment of preferred amino acids. Fine-tuning AlphaFold’s parameters on peptide-MHC binding data does not degrade performance on these other, structurally distinct, classes of protein-peptide interactions; the fine-tuned parameters perform better than the default parameters.

**Figure 4:**
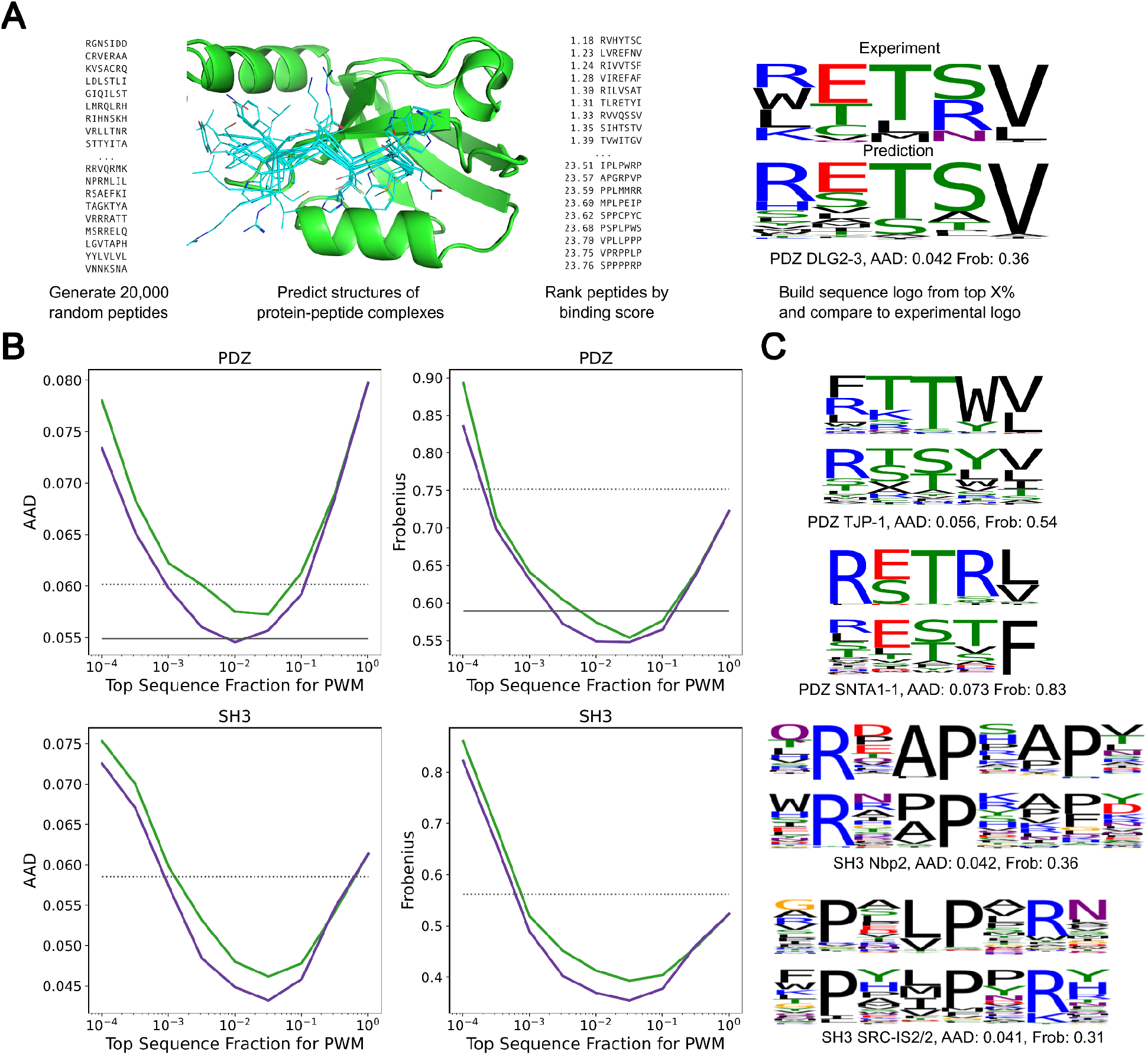
Accurate prediction of PDZ-peptide and SH3-peptide interactions using the combined structure prediction-classification network. A) Prediction of position-weight matrices by random sequence generation followed by network evaluation. B) Combined network fine-tuned on peptide-MHC interactions outperforms AlphaFold with default parameters in recapitulating experimental PWMs. AAD: average absolute difference in amino acid frequency; Frobenius: average Frobenius distance between PWM columns (lower is better for both; see Methods). AlphaFold (green), and fine-tuned AlphaFold (purple). The solid black lines in the PDZ panels indicate the performance of the method from Ref. (16), which was optimized on PDZ domains. Dashed black lines indicate the average AAD and Frobenius scores of PWMs constructed from the sequences of the template PDB peptides used for modeling. C) Examples of predicted and experimental sequence logos (See **Figs. S5-6** for a complete set of sequence logos).

## Discussion

The outstanding performance of AlphaFold on protein structure prediction and NetMHCpan on peptide MHC binding specificity prediction illustrates the power of networks trained on very large datasets; the PDB in the first case, experimental peptide-MHC binding data in the second. Here, we show how to train a single network on both structural and binding data simultaneously, and demonstrate that this enables considerable generalization of binding specificity prediction beyond the available training sets. While sequence-only networks such as NetMHCpan have outstanding performance within their target domains, combined sequence and structure models can be more robust to biological variation and limited training data, and they are able to generalize to new biological systems. Moving forward, there is much to explore in network architectures for combined structure and specificity prediction; our combination of a simple classification module (logistic regression) on top of the AlphaFold network provides a starting benchmark for comparison.

## Materials and Methods

### Peptide-MHC dataset

Structures of peptide-MHC complexes were downloaded from the RCSB Protein Databank (19) with an inclusion cutoff of 2021-08-05. Peptide-MHC binding data was extracted from the training sets generously provided by the NetMHCpan developers on their website (https://services.healthtech.dtu.dk/service.php?NetMHCpan-4.1). The Class I test set in **Figure S4** contained 3236 peptide-MHC pairs (1564 binders and 1672 non-binders) across 98 unique alleles. All examples used for training and testing are provided in the Zenodo repository associated with this manuscript.

### Peptide-MHC template search and query-to-template mapping

The amino acid sequence of the ɑ1 and ɑ2 domains for MHC Class I and the ɑ1 and β1 domains for MHC Class II were considered as the MHC query sequence and were pairwise aligned to their corresponding domains across all entries in the PDB template set. The pairwise alignment was done with the Needleman-Wunsch local alignment algorithm using the BLOSUM62 substitution score matrix while setting gap open penalty to -11 and gap extend penalty to -1. The Biopython package (20) was used to run the alignment algorithm. The alignments were used to generate query-to-template mapping strings that map each query residue to the corresponding residue in the template crystal structure. To generate query-to-template mapping for the peptide, sequence identity was ignored and mappings were generated based on position along the peptide sequence. For Class I molecules, the query peptide was mapped to the template one to one if the length of the two peptides were equal, and if not, the first three and the last three residues of the query were mapped to the corresponding residues of the template. For Class II queries, the core 9mer residues of the peptide and their immediate neighboring residues (11mer residues in total) were mapped to the corresponding residues in the templates. If the 9mer core was not determined in the query, each possible consecutive 11-residue window along the peptide was used as the query peptide to model the complex with all possible positions for the 9mer core and the best model was selected by picking the one with the lowest inter-chain PAE among the windows. Finally, the MHC and peptide mapping strings were combined by putting the peptide either in the N-terminus or the C-terminus of the MHC mapping string.

### AlphaFold modeling of peptide-MHC pairs

Query peptide sequence was appended to the N-terminus or C-terminus of the query MHC sequence. At chain break junctions, the residue indices were shifted by 200. Single sequence and templates were used as inputs and AlphaFold’s model_1, model_1_ptm, model_2, or model_2_ptm weights were used for inference. The crystal structures of the top 4 hits from the template search based on the MHC alignment scores were used to construct the template features in AlphaFold. Both PDB outputs of the predictions and predicted AlphaFold metrics were saved for analysis purposes.

### Hybrid AlphaFold Fine-Tuning on Structure and Binding Data

Combined structure prediction-classification approach for fine-tuning AlphaFold was performed in two stages. First, the binder model parameters were fitted using logistic regression while the AlphaFold parameters were kept fixed at their starting values. The AlphaFold MHC-peptide inter-PAE scores for the training set, together with their binder/non-binder labels, were provided to the `LogisticRegression` class from the Scikit-learn (21) package `linear_model`. This produced slope and intercept values that defined a linear mapping between inter-PAE values and logit values. We noted that these fitted parameters differed between MHC class I and MHC class II: a model fitted on class I peptides had a higher midpoint PAE (8.03) than a model fitted on class II peptides (4.34). This difference likely reflects structural differences in the peptide binding mode, with class I peptides bulging out of the MHC pocket, anchored at the ends, while class II peptides follow a direct and fully extended path with termini protruding at the ends of the pocket. To account for these differences, and allow for future applications in which even more disparate binding data might be combined, we allowed the logistic regression midpoint parameter to vary as a function of a predefined class value (here I or II) for each training example.

The binder model parameters were then fixed at their optimal values while the AlphaFold parameters were fine-tuned in the context of a hybrid structure and binder loss function. The loss for a single training example was equal to the softmax cross entropy of the binder/non-binder prediction for that example plus the AlphaFold structure loss, which was multiplied by a weight of 1.0, if the ground truth structure for that example was an experimentally determined structure, or 0.25, if the ground truth structure was an AlphaFold predicted structure. Predicted structural models were used as the ground truth conformations for the non-binder peptides and for binder peptides without solved structures; for these examples, the AlphaFold structure loss was computed only over the MHC to allow the peptide conformation to adjust during fine-tuning. For the parameters evaluated here, we chose a fairly conservative stopping point after two epochs of training (when each training example had been seen twice) to avoid excessive drift in the AlphaFold model. A Python script that performs parameter fine tuning, together with command line parameters and example inputs, is provided in the github repository associated with this manuscript. Full inputs for fine-tuning, including the predicted AlphaFold models used as ground truth structures, are provided in the Zenodo repository associated with this manuscript.

### Evaluation of classification performance on peptide-MHC

Pairs of peptide-MHC from the dataset under evaluation were modeled with AlphaFold or Fine-Tuned AlphaFold parameters. Inter-chain PAE terms corresponding to the peptide-MHC interactions were averaged. Mean inter-chain PAE values and binder non-binder class labels were used to construct Receiver Operating Characteristic (ROC) curves and the area under the curve values were used to compare classification performance of different models. To compare our prediction results to NetMHCpan, we used NetMHCpan-4.1 and NetMHCIIpan-4.1 (1, 2). These software tools were installed on our server and executed through the command line.

### AlphaFold modeling of PDZ and SH3 domains

The PDZ domain dataset (**Table S1**) was taken from Ref. (16) and consisted of 17 human PDZ domains with experimentally determined structures. Binder peptides for the 17 PDZ domains were downloaded from supplemental data for Ref. (17) (https://baderlab.org/Data/PDZ). Experimental amino acid frequency matrices (PWMs) were constructed from the PDZ binder peptides, with clone frequency weighting. For the AlphaFold simulations, 20,000 random peptide sequences of length equal to the peptide in the experimental structure were generated using NNK codon frequencies to match the amino acid bias in the phage display libraries. The experimental structure listed in the ‘template’ column of **Table S1** was used as the sole template, with the random peptide sequences aligned to the template peptide and single-sequence MSA information.

The SH3 domain dataset (**Table S2**) consisted of 19 SH3 domains with experimentally determined structures extracted from the Database of Peptide Recognition Modules (http://prm-db.org/) (18). Experimental PWMs were downloaded from the PRM-DB. SH3 domains can bind peptides in two orientations, denoted class I and class II, which have opposite chain orientations: class I peptides often match a +XXPXXP sequence motif, where ‘+’ denotes a positively charged amino acid and X is any amino acid; class II peptides often match a PXXPX+ motif. SH3 peptide PWMs from PRM-DB were annotated as class I or class II by choosing the class whose sequence motif had the highest PWM frequency (averaged over the three motif positions). Five of the SH3 domains had multiple PWMs in the PRM-DB, one of which was assigned as class I and one as class II; these domains were modeled twice, once in each orientation. For AlphaFold modeling, the native PDB structure listed in **Table S2** (‘SH3 template’ column) was used as the template for the SH3 domain. Four peptide-SH3 structures with peptides in the desired orientation (i.e., class I or class II) were chosen as peptide templates based on SH3 domain sequence identity (**Table S2**, ‘Peptide templates’ column). The peptides in these structures were transformed into the reference frame of the SH3 domain template by structural superimposition to create hybrid template models for AlphaFold. Multiple structural alignment was used to identify the core motif positions (+XXPXXP or PXXPX+) in each template peptide. The peptide sequence modeled in the AlphaFold runs consisted of the core motif together with one residue on either side (9 residues for class I and 8 residues for class II).

For comparison with experimental PWMs, predicted PWMs were constructed from the top ranked peptide sequences. Peptides were ranked by protein-peptide inter-PAE: the sum of the residue-residue PAE scores for all (protein,peptide) and (peptide,protein) residue pairs, where PAE is AlphaFold’s ‘predicted aligned error’ accuracy measure. The experimental PWMs were derived from phage display experiments with random peptide libraries of size 10^9^ and greater, whereas the predicted PWMs were based on 20,000 modeled peptides. To account for this differential and better match the entropy of the amino acid frequency distributions, we squared the predicted amino acid frequencies and renormalized them to sum to 1. This had the effect of increasing the information content of the predicted PWMs without changing the order of amino acid preference. The exponent of 2 can be partly rationalized by the approximate two-fold differential in log search-space size between predictions and experiments. Following Ref. (16), predicted and experimental PWMs for PDZ domains were compared over the last 5 C-terminal peptide positions. Predicted and experimental PWMs for SH3 domains were compared over the core 7 (for class I) or 6 (for class II) positions of the SH3 motif together with the immediately adjacent positions, if those positions were present in the experimental PWM. Two measures of PWM column divergence were used to assess predictions: average absolute deviation (AAD) and the Frobenius metric. The AAD for a single PWM position equals the sum of the absolute frequency deviations for all amino acids, divided by 20; AAD ranges from 0.0 (perfect agreement) to 0.1 (maximal divergence). The Frobenius metric for a single PWM position equals the square root of the sum of the squared frequency deviations; it ranges from 0.0 (perfect agreement) to the square root of 2 (maximal divergence). The AAD and Frobenius values in **Figure 4** were averaged over all PWM columns.

### HCV experiment

We scanned the sequence of the Hepatitis C Virus (HCV) genome polyprotein (3011 amino acids; Uniprot ID P27958) to find potential 10mer peptide epitopes presented by HLA-A*02:01. We ran NetMHCpan with FASTA format input and default parameters. NetMHCpan compares 10mer peptides to its internal 9mer binding model by dropping one “gap” position from the peptide:model alignment. We parsed the location of the gap position from the output columns (‘Of’, ‘Gp’, and ‘Gl’) and confirmed by matching to the sequence in the ‘Core’ column. To assess divergence between our structure-based predictions and NetMHCpan, raw scores from each method were sorted and converted to rank scores. Peptides were grouped by the NetMHCpan-assigned gap position, and the mean absolute difference in rank score was computed for each gap position.

## Acknowledgements

The authors wish to thank Frank DiMaio, Sergey Ovchinnikov, Ivan Anishchanka, Amijai Saragovi, Aditya Krishnakumar, and Ian Humphreys for helpful discussions, Morten Nielsen for assistance with NetMHCpan, and Luki Goldschmidt for technical IT support. We are grateful to the developers of NetMHCpan for making their training and testing datasets publicly available.

## Funding

We thank AWS and Microsoft for providing computing resources. This work was supported with funds provided by a Microsoft gift (A.M., J.D., M.B., D.B.); the Audacious Project at the Institute for Protein Design (D.B.); the Howard Hughes Medical Institute (M.A., D.B.); and NIH R35 GM141457 (P.B.). The Jane Coffin Childs Memorial Fund for Medical Research (M.A).

## Author contributions

Conceptualization: P.B., D.B.

Methodology: A.M., P.B., J.D.

Software: A.M., P.B., J.D., M.B.

Validation: A.M., P.B.

Formal analysis: A.M., P.B., J.D, M.A.

Resources: P.B., D.B.

Data curation: P.B.

Writing–original draft: A.M., P.B., D.B.

Writing–review & editing: A.M., P.B., D.B., J.D., M.B., M.A.

Visualization: A.M., P.B.

Supervision: P.B., D.B.

Project administration: A.M., P.B., D.B.

Funding acquisition: P.B., D.B.

## Competing interests

Authors declare that they have no competing interests.

## Data and code availability

Fine tuning the AlphaFold model parameters required small changes to the AlphaFold software. The modified version of the AlphaFold package used here as well as additional python scripts for training and prediction will be made available prior to publication at https://github.com/phbradley/alphafold_finetune. Full datasets for training and testing the structure prediction-classification model, including predicted AlphaFold structures for binder and non-binder examples, will be made available prior to publication in the Zenodo repository <insert repository ID>. [This preprint text will be updated when these resources are released].

## Supplementary Material

**Supplementary Table 1.**
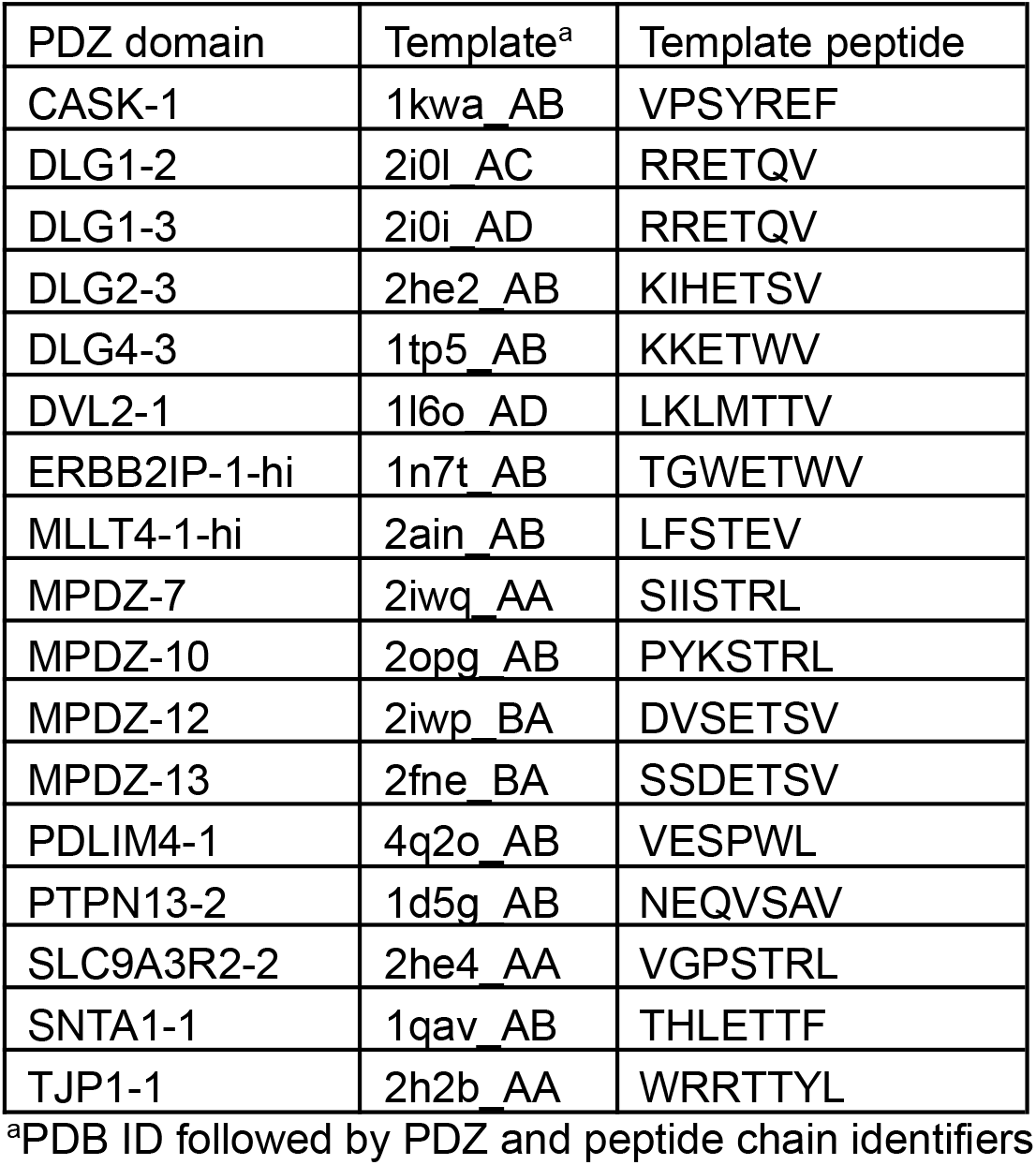
PDZ domain dataset.

**Supplementary Table 2.** SH3 domain dataset.

| PRM-DB identifier<sup>a</sup> | SH3 class | Protein name | Uniprot identifier | SH3 template<sup>b</sup> | Peptide templates<sup>c</sup> |
| --- | --- | --- | --- | --- | --- |
| PRM_0152 | 1 | Sho1 | P40073:287-367 | 2vkn_A | 2vkn_AC,3ua7_AE,4rtz_AB,5sxp_BF |
| PRM_0157 | 1 | GRB2 | P62993:1-58 | 1gbq_A | 5sxp_BF,1io6_AB,2bz8_AC,3ua7_AE |
| PRM_0161 | 2 | Pex13 | P80667:295-386 | 1n5z_A | 1n5z_AP,1gbq_AB,1gbq_AB,5ul6_AM |
| PRM_0162 | 1 | Nbp2 | Q12163:103-180 | 2lcs_A | 2lcs_AB,2bz8_AC,1io6_AB,4rtz_AB |
| PRM_0166 | 2 | Lsb3 | P43603:385-451 | 1ssh_A | 1ssh_AB,3ua7_DF,2d1x_DQ,5ul6_AM |
| PRM_0190 | 2 | Bbc1 | P47068:1-78 | 1zuk_B | 2rqu_AB,4ln2_AB,2d1x_DQ,2rpn_AB |
| PRM_0193 | 2 | Abp1 | P15891:535-592 | 2rpn_A | 2rpn_AB,2d1x_DQ,4wci_EF,2rqu_AB |
| PRM_0210 | 1 | GRB2 | P62993:1-57 | 1gbq_A | 5sxp_BF,1io6_AB,2bz8_AC,3ua7_AE |
| PRM_0210 | 2 | GRB2 | P62993:1-57 | 1gbq_A | 1gbq_AB,1gbq_AB,2d1x_DQ,5xhz_AC |
| PRM_0215 | 1 | SRC-IS2/2 | P12931:87-144 | 4rtz_A | 4rtz_AB,3ua7_AE,5sxp_BF,1io6_AB |
| PRM_0215 | 2 | SRC-IS2/2 | P12931:87-144 | 4rty_A | 4rty_AB,3ua7_DF,5ul6_AM,4f14_AB |
| PRM_0231 | 2 | ASAP1 | Q9ULH1:1073-1131 | 2rqu_A | 2rqu_AB,5xhz_AC,2rpn_AB,2d1x_DQ |
| PRM_0238 | 2 | ARHGEF7 | Q14155:187-242 | 5sxp_B | 5xhz_AC,3u23_AB,2d1x_DQ,3ua7_DF |
| PRM_0266 | 1 | FYN | P06241:85-142 | 3ua7_A | 3ua7_AE,4rtz_AB,5sxp_BF,1io6_AB |
| PRM_0266 | 2 | FYN | P06241:85-142 | 3ua7_D | 3ua7_DF,4rty_AB,5ul6_AM,1gbq_AB |
| PRM_0270 | 2 | CRK | P46108:135-191 | 5ul6_A | 5ul6_AM,4rty_AB,1gbq_AB,1gbq_AB |
| PRM_0272 | 2 | CD2AP | Q9Y5K6:111-166 | 3u23_A | 3u23_AB,5xhz_AC,4wci_EF,2d1x_DQ |
| PRM_0273 | 2 | CD2AP | Q9Y5K6:2-58 | 4wci_E | 4wci_EF,5xhz_AC,3u23_AB,2rpn_AB |
| PRM_0282 | 2 | CTTN | Q14247:459-512 | 2d1x_D | 2d1x_DQ,4f14_AB,2rpn_AB,1gbq_AB |
| PRM_0285 | 1 | SH3KBP1 | Q96B97:2-57 | 2bz8_A | 2bz8_AC,5sxp_BF,1io6_AB,3ua7_AE |
| PRM_0285 | 2 | SH3KBP1 | Q96B97:2-57 | 2bz8_A | 4wci_EF,2d1x_DQ,5xhz_AC,3u23_AB |
| PRM_0291 | 1 | BIN1 | O00499:523-593 | 5i22_A | 5sxp_BF,2bz8_AC,1io6_AB,3ua7_AE |
| PRM_0298 | 1 | SORBS1 | Q9BX66:870-927 | 4ln2_A | 5sxp_BF,2bz8_AC,1io6_AB,3ua7_AE |
| PRM_0298 | 2 | SORBS1 | Q9BX66:870-927 | 4ln2_A | 4ln2_AB,1gbq_AB,1gbq_AB,2rqu_AB |

<sup>a</sup>Internal identifier used in the PRM-DB database (http://prm-db.org/)
<sup>b</sup>PDB and chain identifier of SH3 domain modeling template
<sup>c</sup>PDB identifier followed by SH3 and peptide chain identifiers of peptide modeling templates

**Supplementary Figure 1.**
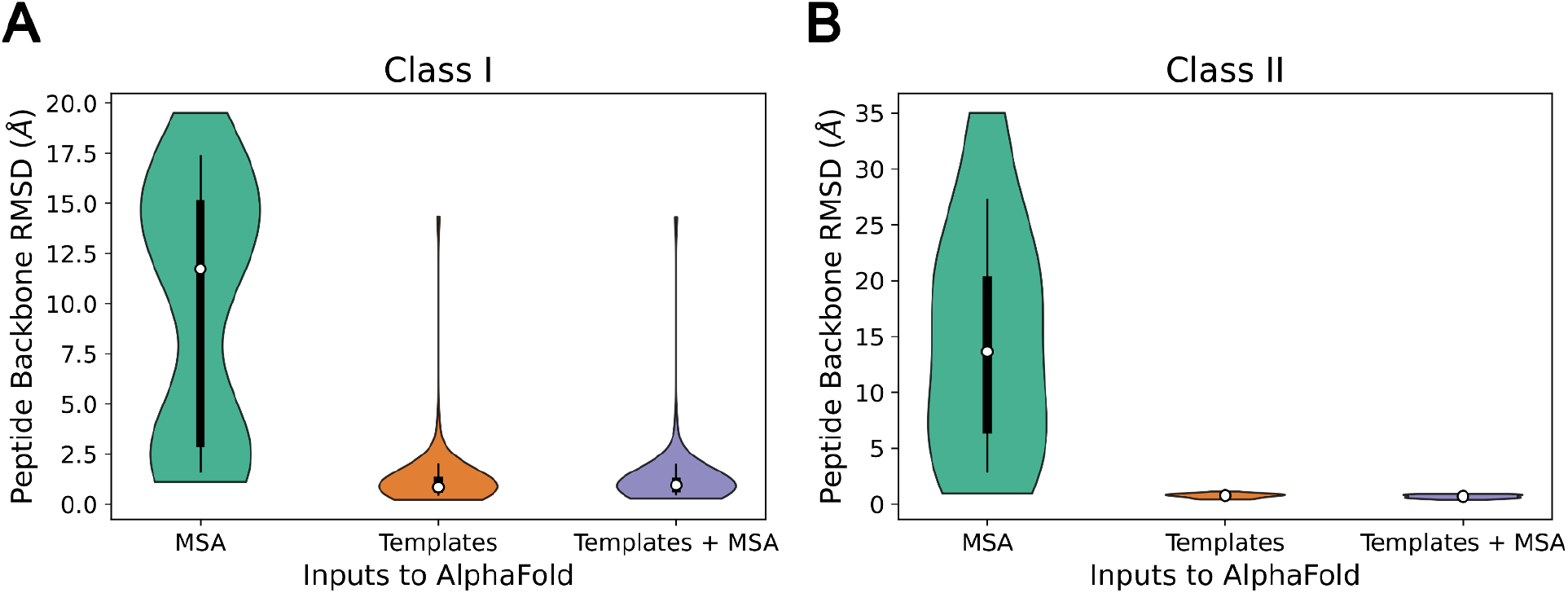
Comparison of peptide-MHC structure modeling quality of AlphaFold with different inputs. **A, B**. Violin plots of peptide backbone RMSD distribution of peptide-MHC models produced by AlphaFold with MSA, Templates, or MSA and Templates as inputs to the network for (A) Class I and (B) Class II complexes. White circles represent the median, thick lines the interquartile range, and thin lines the range between 10th and 90th percentiles.

**Supplementary Figure 2.**
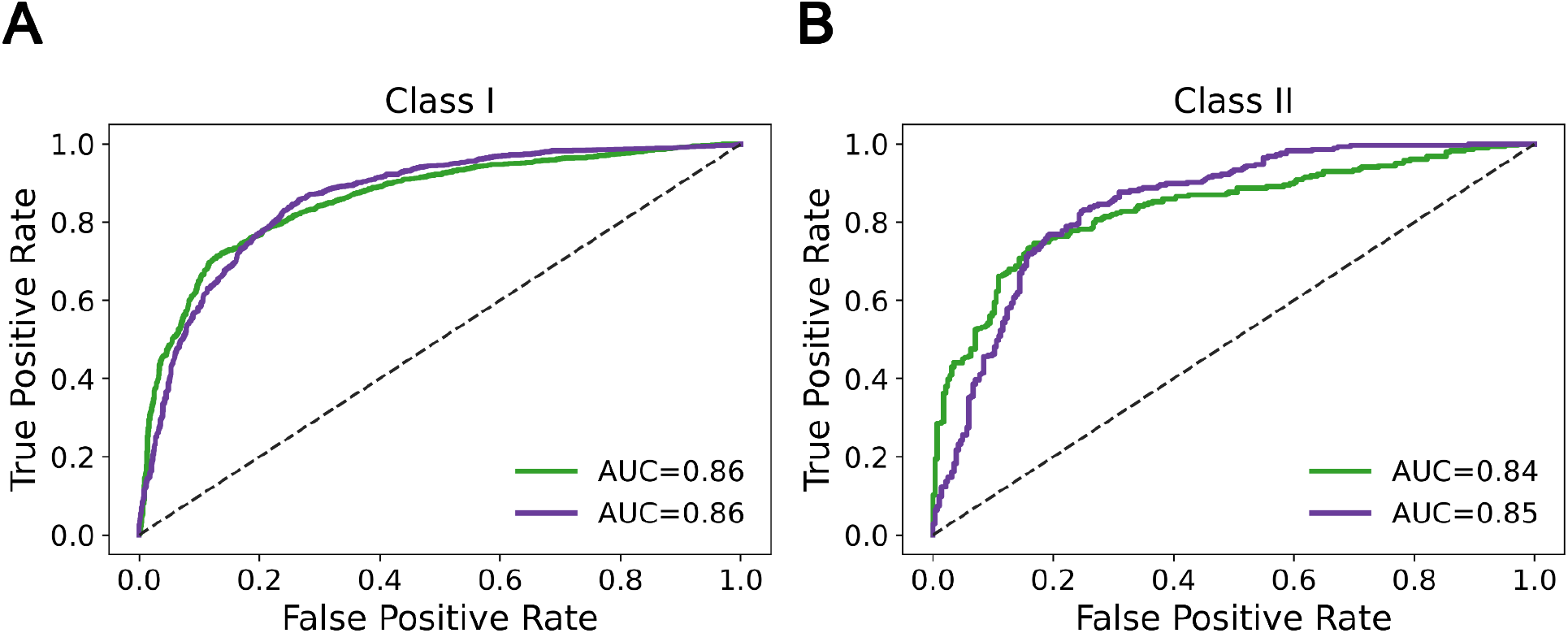
Peptide-MHC classification with AlphaFold structure accuracy metrics. **A, B**. ROC curves comparing AlphaFold’s confidence metrics in binder/non-binder peptide classification for (A) MHC Class I and (B) MHC Class II. Mean inter-chain PAE (green) and mean peptide pLDDT (purple).

**Supplementary Figure 3.**
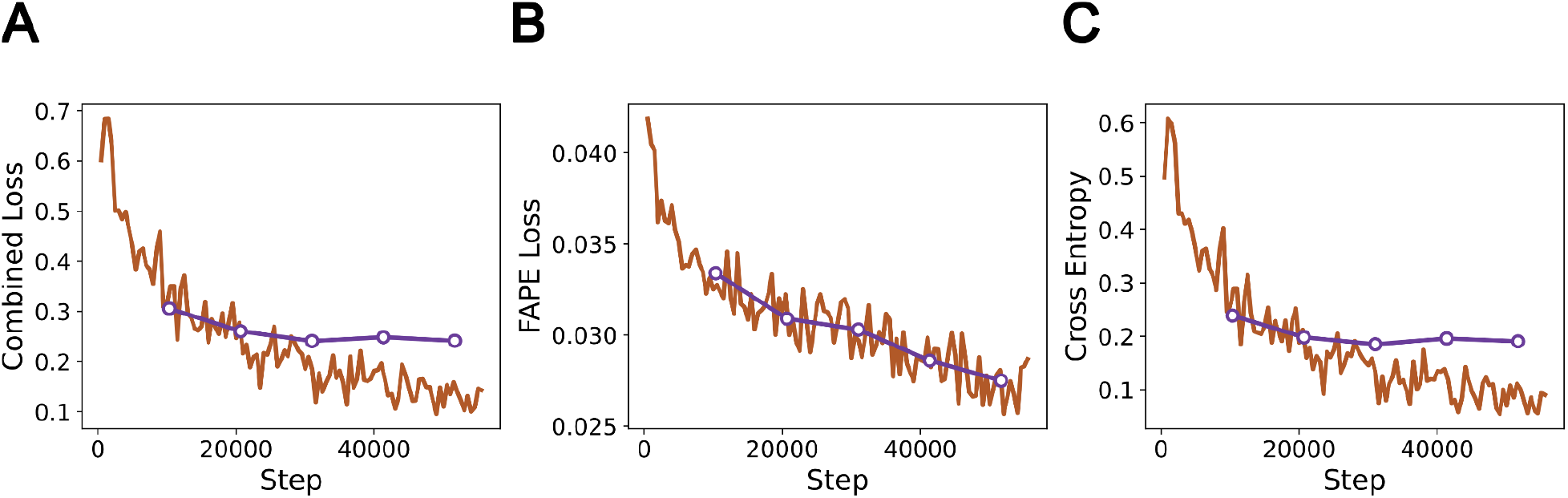
Loss progression curve during fine-tuning of AlphaFold on peptide-MHC structures. **A-C**. Plots of (A) combined fine-tuning loss, (B) FAPE loss on structures (crystal structures and predicted self-distillation structures), and (C) cross entropy of peptide-MHC binding classification against training steps in the combined structural and classification fine-tuning of AlphaFold parameters on peptide-MHC data. Training loss (maroon) and validation loss (purple).

**Supplementary Figure 4:**
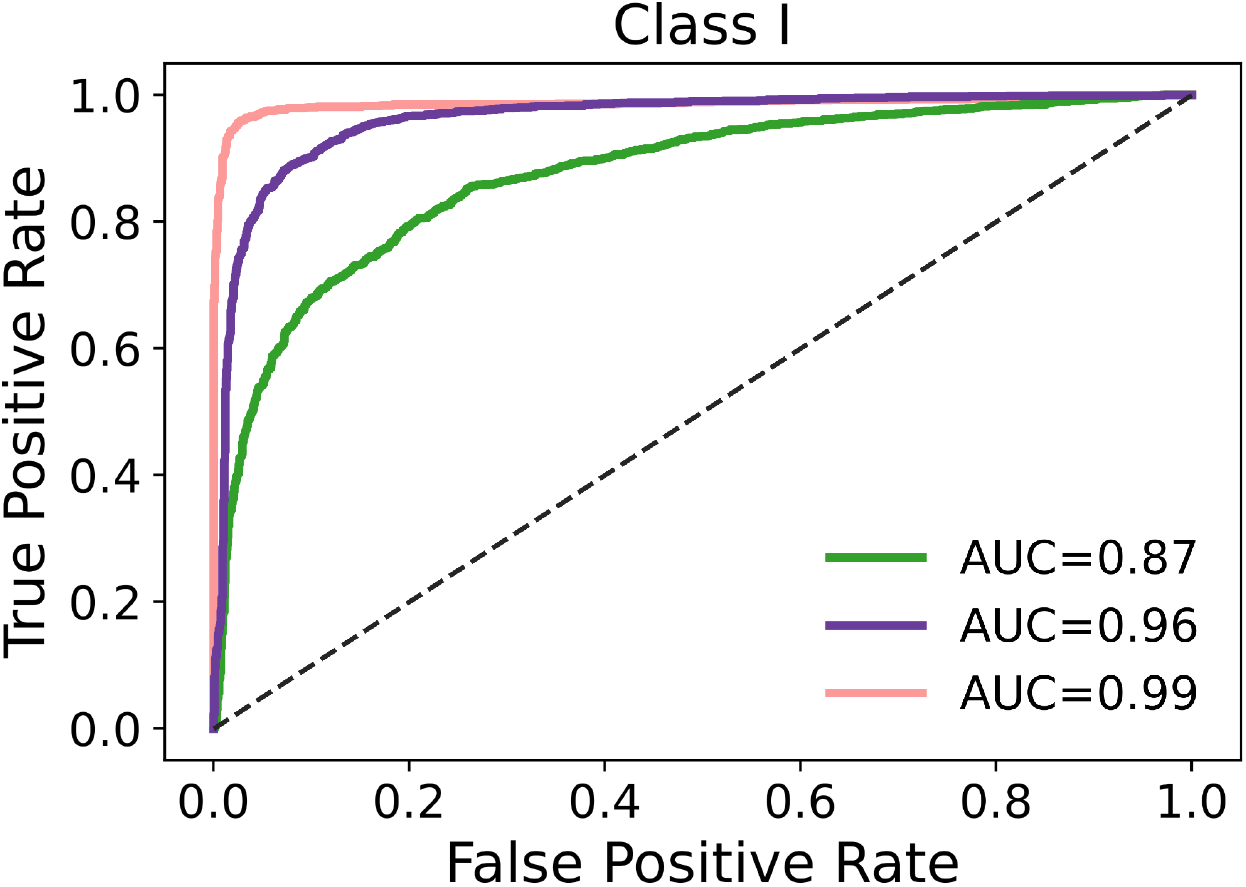
Classification performance with Class I test set excluding similarly-labeled peptides in common with the training set. We evaluated binder/non-binder classification performance of the fine-tuned model on a separate test set filtered to exclude peptides that always occur with the same label in both the train and test set. In the training of NetMHCpan, positive peptides overlapping with the training set were excluded from the test set; here we exclude identical peptides when all their occurrences in the training and test set are of the same label. AlphaFold (green), combined structure prediction-classification model (purple), and NetMHCpan (pink).

**Supplementary Figure 5.**
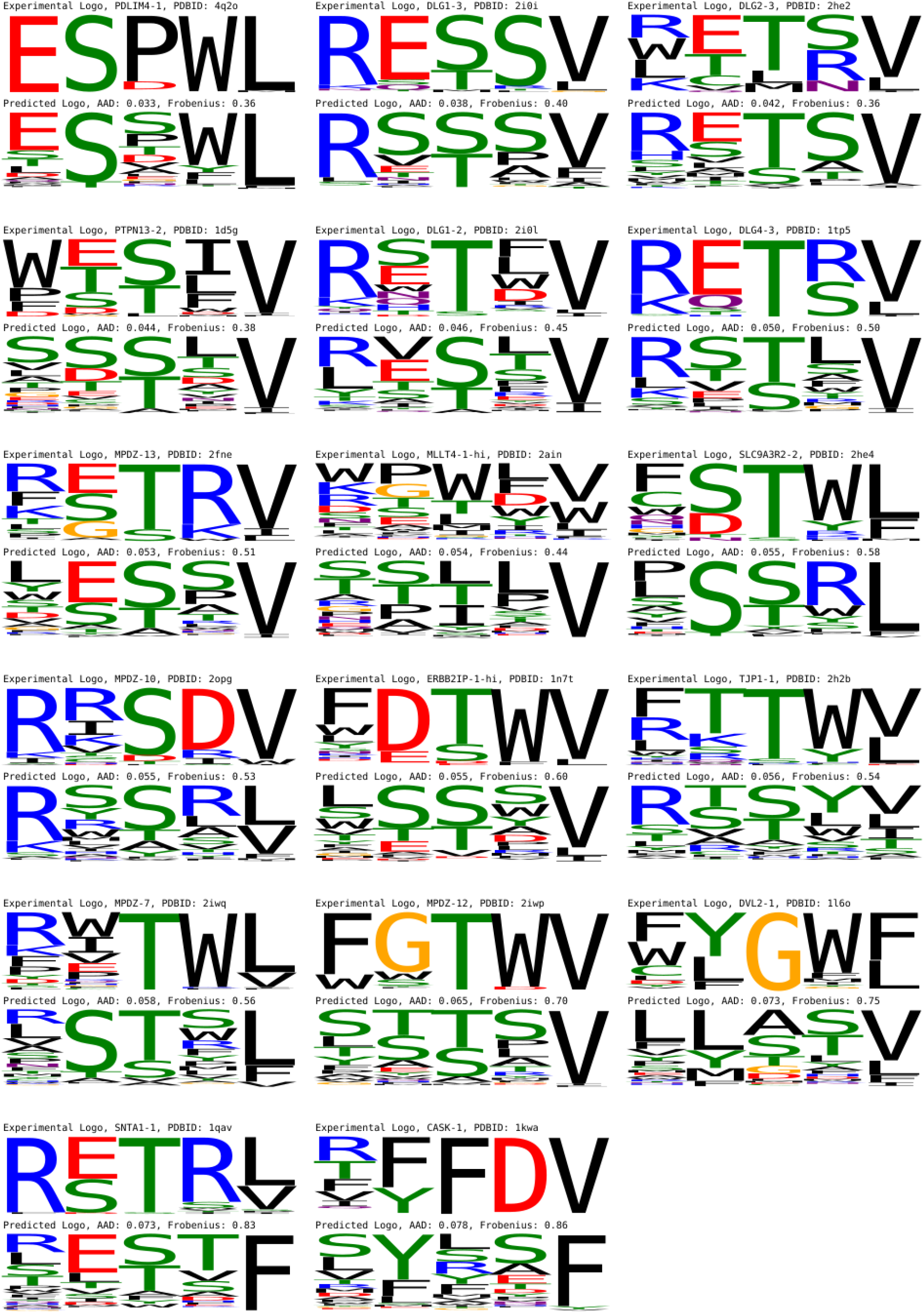
Predicted and experimental sequence logos for the 17 PDZ domains. Predicted logos were built from the top 1% of random peptides ranked by the combined structure prediction-classification model. Domains are ordered by increasing AAD between predicted and experimental logos.

**Supplementary Figure 6.**
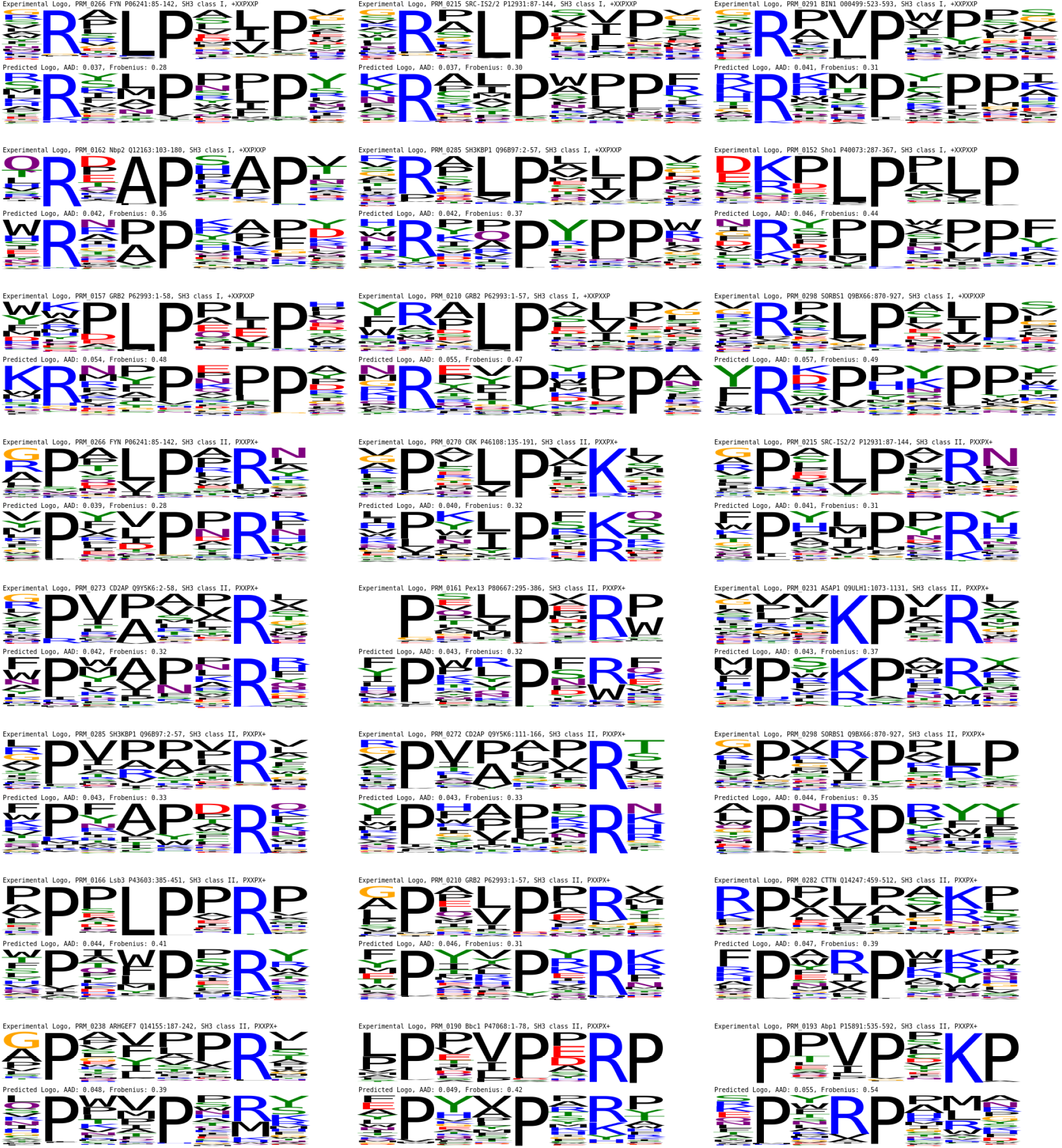
Predicted and experimental sequence logos for the 24 SH3 domains. Predicted logos were built from the top 1% of random peptides ranked by the combined structure prediction-classification model. Domains are grouped by SH3 class and ordered within each class by increasing AAD between predicted and experimental logos.

## Notes

https://github.com/phbradley/alphafold_finetune

### Competing Interest Statement

The authors have declared no competing interest.

